# Freeze-thaw Cycles Enable a Prebiotically Plausible and Continuous Pathway from Nucleotide Activation to Nonenzymatic RNA Copying

**DOI:** 10.1101/2021.09.07.459201

**Authors:** Stephanie J. Zhang, Daniel Duzdevich, Christopher E. Carr, Jack W. Szostak

**Affiliations:** Department of Chemistry and Chemical Biology, Harvard University, 12 Oxford Street, Cambridge, Massachusetts 02138, United States; Department of Molecular Biology, Center for Computational and Integrative Biology, Massachusetts General Hospital, Boston, MA 02114, United States of America; Howard Hughes Medical Institute, Massachusetts General Hospital, Boston, MA 02114, United States of America; Daniel Guggenheim School of Aerospace Engineering, School of Earth and Atmospheric Sciences, Georgia Institute of Technology, Atlanta, GA 30332, United States of America

**Keywords:** Chemical nucleotide activation, RNA copying, Origin of life, RNA world hypothesis, Prebiotic chemistry

## Abstract

Nonenzymatic template-directed RNA copying using chemically activated nucleotides is thought to have played a key role in the emergence of genetic information on the early Earth. A longstanding question concerns the number and nature of different environments that might have been necessary to enable all of the steps from nucleotide synthesis to RNA replication. Here we explore three sequential steps from this overall pathway: nucleotide activation, synthesis of imidazolium-bridged dinucleotides, and template-directed primer extension. We find that all three steps can take place in one reaction mixture, under conditions of multiple freeze-thaw cycles. Recent experiments have demonstrated a potentially prebiotic methyl isocyanide-based nucleotide activation chemistry. Unfortunately, the original version of this approach is incompatible with nonenzymatic RNA copying because the high required concentration of the imidazole activating group prevents the accumulation of the essential imidazolium-bridged dinucleotide needed for primer extension. Here we report that ice eutectic phase conditions facilitate not only the methyl isocyanide-based activation of ribonucleotide 5′-monophosphates with stoichiometric 2-aminoimidazole, but also the subsequent conversion of these activated mononucleotides into imidazolium-bridged dinucleotides. Furthermore, this one pot approach is compatible with template-directed primer extension in the same reaction mixture. Our results suggest that the simple and common environmental fluctuation of freeze-thaw cycles could have played an important role in prebiotic nucleotide activation and nonenzymatic RNA copying.

**Significance Statement:** The replication of RNA without the aid of evolved enzymes may have enabled the inheritance of useful molecular functions during the origin of life. Several key steps on the path to RNA replication have been studied in isolation, including chemical nucleotide activation, synthesis of a key reactive intermediate, and nonenzymatic RNA copying. Here we report a prebiotically plausible scenario under which these reactions can happen together under mutually compatible conditions. Thus, this pathway could potentially have operated in nature without the complicating requirement for exchange of materials between distinct environments.

## Introduction

Template-directed nonenzymatic RNA copying requires activated nucleotides. A range of phosphate-activating groups have been explored to chemically activate nucleotides for such polymerization reactions.^1-2^ Our laboratory has demonstrated efficient copying of several short RNA templates using 2-aminoimidazole (2AI) activated ribonucleotides (2AImpN, *pN),^3^ and shown that polymerization proceeds predominantly through spontaneously generated 5′-5′-imidazolium-bridged dinucleotides.^4^

Sutherland and co-workers recently reported a potentially prebiotic route to selective phosphate activation with methyl isocyanide (MeNC), aldehyde, and imidazole.^5^ MeNC could have been produced in a ferrocyanide- and nitroprusside-containing environment upon UV irradiation.^5^ A major hurdle to the compatibility of this activation chemistry with RNA copying is that activation requires an excess of 2AI to drive the reaction forward; however, excess 2AI prevents nonenzymatic RNA polymerization by inhibiting the accumulation of imidazolium-bridged dinucleotides.^4, 6^ We have identified a variant of isocyanide activation chemistry, named bridge-forming activation, that circumvents this issue by directly generating bridged dinucleotides from activated mononucleotides.^7^ The resulting high concentration of bridged dinucleotides increases the yield of primer extension products.

Although we were able to demonstrate the compatibility of bridge-forming activation with nonenzymatic RNA polymerization, bridge-forming chemistry depends on the presence of previously activated mononucleotides and the absence of excess 2AI. To begin to address the challenge of initial nucleotide activation under conditions that do not inhibit bridged dinucleotide accumulation, we have explored the effects of repeated freeze-thaw cycles on activation chemistry.^8-9^

Under partially frozen conditions, solutes are excluded from growing ice crystals and become concentrated in the interstitial liquid solution. This phenomenon of eutectic phase concentration has been demonstrated to facilitate the synthesis of nucleotide precursors,^10^ as well as the condensation of activated nucleotides into oligomers.^11^ Here we report the *in situ* activation of mononucleotides with stoichiometric 2AI by repeatedly subjecting the solution to eutectic phase freezing, which temporarily brings the reactants into close physical proximity and obviates the need for the large excess of 2AI otherwise required to drive activation in relatively dilute liquid phase reactions. We find that all four canonical ribonucleotides can be activated with this approach, which also yields high concentrations of bridged dinucleotides. Consequently, we were also able to show that the reaction is compatible with template-directed nonenzymatic RNA copying. We then apply a deep-sequencing assay to demonstrate that the high concentrations of bridged dinucleotides generated by ice eutectic phase activation promote relatively high-fidelity copying reactions. We also show that in a one-pot reaction, the ice eutectic phase does not induce an excess of mismatches during primer extension. The geochemically plausible conditions that we have identified enable a potentially prebiotic pathway that begins with ribonucleotide 5′-monophosphates, generates 2AI activated nucleotides and then the essential bridged dinucleotides, and finally yields template-directed primer extension products, all in one reaction mixture.

## Results

Addressing the incompatibility between template-directed primer extension and the initial *in situ* activation of mononucleotides requires a plausible pathway to nucleotide activation using a concentration of 2AI comparable to the concentration of nucleotides, rather than the large excess currently required (Figure 1A). We reasoned that the ice eutectic phase as a medium for activation could temporarily increase the effective local concentration of 2AI, thereby alleviating the requirement for an absolute excess of 2AI relative to nucleotides (Figure 1B). To begin exploring the effect of ice eutectic phase concentration on nucleotide activation, we treated 10 mM adenosine monophosphate (pA) with 10 mM 2AI, 100 mM 2-methylbutyraldehyde (2MBA), and 200 mM MeNC, at pH 8 under both room temperature and ice eutectic phase conditions.^7^ With these reactant concentrations, and incubation at room temperature, only 2.0±0.1 mM activated nucleotides (approximately 20% of the 10 mM total available nucleotides) and 0.11±0.01 mM bridged dinucleotides were formed after 24 hours (Figure 1C). In contrast, using the same concentrations of reactants but subjecting the mixture to ice eutectic phase concentration at -13 °C for 24 hours yielded 50% total activation, consisting of 3.0±0.1 mM activated nucleotides and 1.0±0.1 mM bridged dinucleotides. The ice eutectic phase not only increases the yield of activated nucleotides but also acts as a storage mechanism, because activated nucleotides are less susceptible to hydrolysis at low temperatures (Figure S1C-D), which further aids in the accumulation of bridged dinucleotides (Figure 1C).

**Figure 1.**
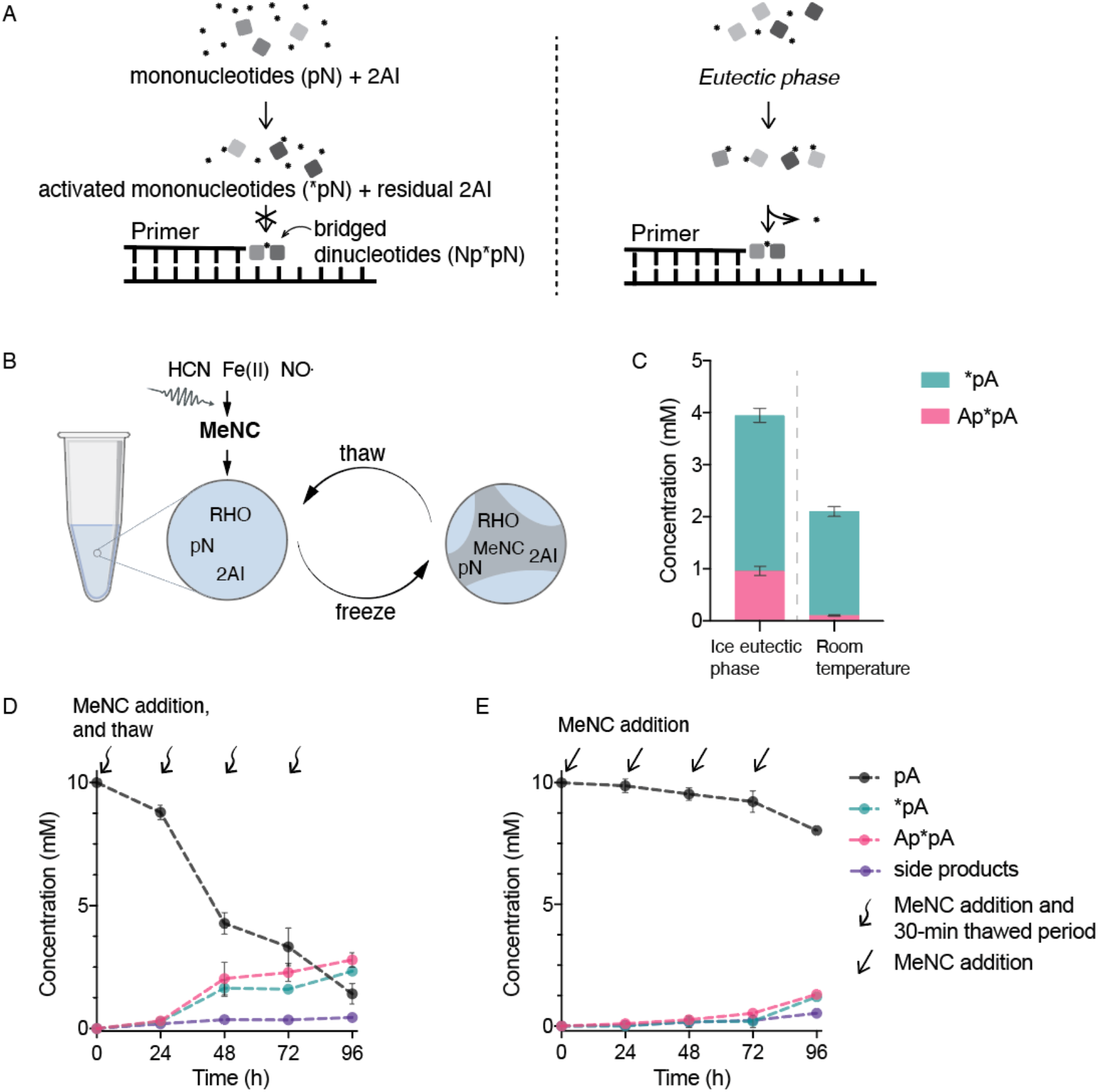
Ice eutectic phase significantly enhances the yield of activated nucleotides (*pN) and promotes formation of bridged dinucleotides (Np*pN). (A) Scheme showing that ice eutectic concentration enables monophosphate ribonucleotide activation and template-directed RNA copying in one reaction. At room temperature (left panel), the excess 2AI needed to drive nucleotide activation also inhibits bridged dinucleotide accumulation and therefore template copying. In contrast, using eutectic ice during activation (right panel) requires only equimolar concentrations of monophosphate ribonucleotide and 2AI, allowing bridged dinucleotide accumulation and template copying. (B) Scheme of how eutectic ice concentrates solutes in the liquid phase in between the pure-water ice crystals. (C) Effect of ice eutectic phase on nucleotide activation. Nucleotide activation at ice eutectic phase (left) and room temperature (right) with one round of MeNC addition and equimolar concentrations of initially unactivated nucleotides and 2AI. (The complete breakdown of side-products is included in Figure S3A.) (D) Cycles of MeNC addition and eutectic freezing drive significant nucleotide activation. Controlled addition of MeNC to the thawed solution enhances the yield of activated nucleotides (*pA). The curvy arrows indicate the addition of MeNC after thawing of the reaction mixture. Note that the reaction mixture is frozen as eutectic ice for the entire reaction course except during the brief thawing steps (∼30 minutes at room temperature for the mixture to thaw completely). (The complete breakdown of side-products is included in Figure S3B.) (E) Activation is inefficient in the absence of ice eutectic freezing, at room temperature. The straight arrows indicate the addition of MeNC at room temperature. (The complete breakdown of side-products is included in Figure S3C.) *Reaction conditions*: 10 mM pA, 10 mM 2AI, 100 mM 2MBA, 30 mM MgCl_2_, 50 mM Na^+^-HEPES pH 8, plus (C) one addition of 200 mM MeNC at ice eutectic phase or room temperature, or (D) periodic addition of the MeNC beyond the initial 50 mM in three aliquots of 50 mM each under ice eutectic phase at -13 °C or (E) at room temperature. Errors are standard deviations of the mean, n ≥ 2 replicates.

The possibility that in a prebiotic environment MeNC could have been released periodically in response to day/night cycles of UV irradiation^12^ prompted us to test multiple cycles of eutectic phase concentration and MeNC delivery. Instead of adding 200 mM MeNC from the start, 50 mM MeNC was introduced to the solution at the beginning of each of four sequential freeze-thaw cycles. This approach further increased nucleotide activation to 75%, yielding 2.33±0.01 mM activated nucleotides and 2.8±0.3 mM bridged dinucleotides (Figure 1D). By comparison, periodic addition of MeNC at room temperature without eutectic freezing yielded only 1.30±0.6 mM activated nucleotides and 0.7± 0.6 mM bridged dinucleotides, or 27% total activation (Figure 1E). A range of pHs was examined to rule out the possibility that freezing-induced changes in pH might account for the increase in activation (Figure S2).

We then varied the frequency of MeNC addition to the system and evaluated the effect on activated nucleotide yield. We found that an interval of roughly one day between MeNC additions led to optimal activation (Figure 2A-B). Further increases in the interval, up to several days, did not significantly change the activation yield (Figures 2 & S2 (B) to (D)). This not only indicates the robustness of the system in response to changes in the timing of activation but again demonstrates that freezing is a storage mechanism in our system: the accumulation of bridged dinucleotides is consistent with the expectation that activated mononucleotides and bridged dinucleotides should not readily hydrolyze in the ice eutectic state (Figure S1).^6, 13^ In addition, the presence of high concentrations of bridged dinucleotides suggests that bridge-forming chemistry is also occurring. To test this, we incubated 10 mM pure activated mononucleotides in the ice eutectic phase without additional activation chemistry and observed only 1.05±0.05 mM bridged dinucleotides (Figure S4) compared to 2.8±0.3 mM bridged dinucleotides in the presence of activation reagents. Thus, bridge-forming chemistry proceeds under ice eutectic phase conditions.

**Figure 2.**
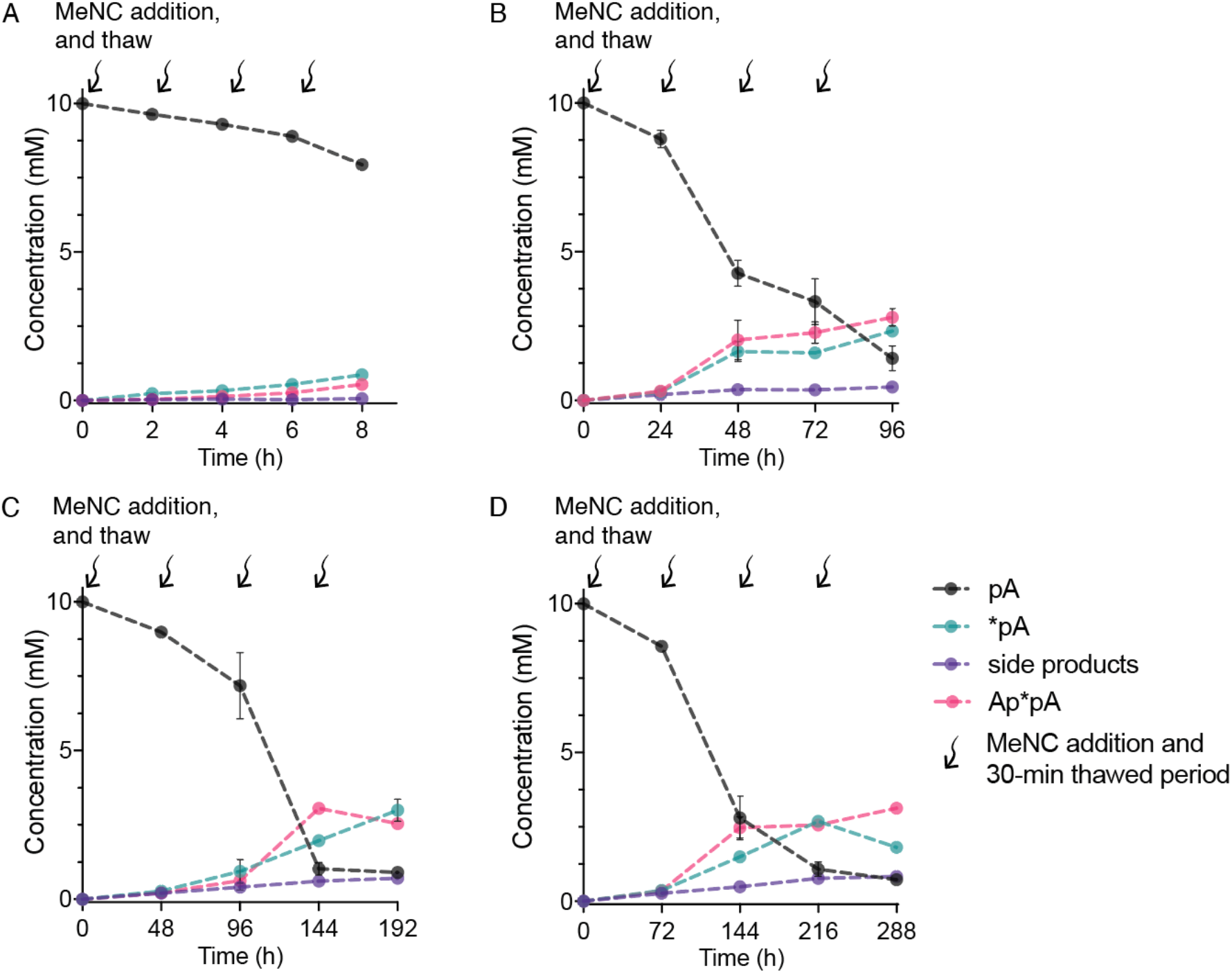
The frequency of MeNC delivery at long time intervals does not affect the overall yield of activated products. (A) Yields of activated species are low with time intervals on the order of hours in between MeNC additions. (B-D) However, lengthening of the time interval beyond 24 hours does not significantly affect the final yield. The complete product breakdown is included in Figure S5. *Reaction conditions*: 10 mM pA, 10 mM 2AI, 100 mM 2MBA, 50 mM MeNC, 30 mM MgCl_2_, 50 mM Na^+^-HEPES pH 8, and subsequent periodic addition of MeNC in three aliquots of 50 mM. Errors are standard deviations of the mean, n ≥ 2 replicates.

Copying mixed sequence RNA templates would require G, C, and U in addition to A. We therefore proceeded to test the activation of all four canonical nucleotides, separately and together, under ice eutectic phase conditions. We began with 10 mM of each individual ribonucleotide and found good yields of activated species with cytidine monophosphate (pC) and uridine monophosphate (pU). In contrast, guanosine monophosphate (pG) exhibited poor activation (Figure 3A). We were able to rescue high efficiency activation with minimal side product formation by treating a mixture of all four nucleotides (10 mM total) with activation chemistry under ice eutectic conditions (Figure 3B). Dilution experiments showed that the rescue is partially due to the decreased relative concentration of pG (from 10 mM to 2.5 mM, Figure 3C), suggesting that concentration-driven aggregation may interfere with the activation of pG. Our observation is another striking example of the desirability of an optimal degree of heterogeneity in prebiotic processes.^14^

**Figure 3.**
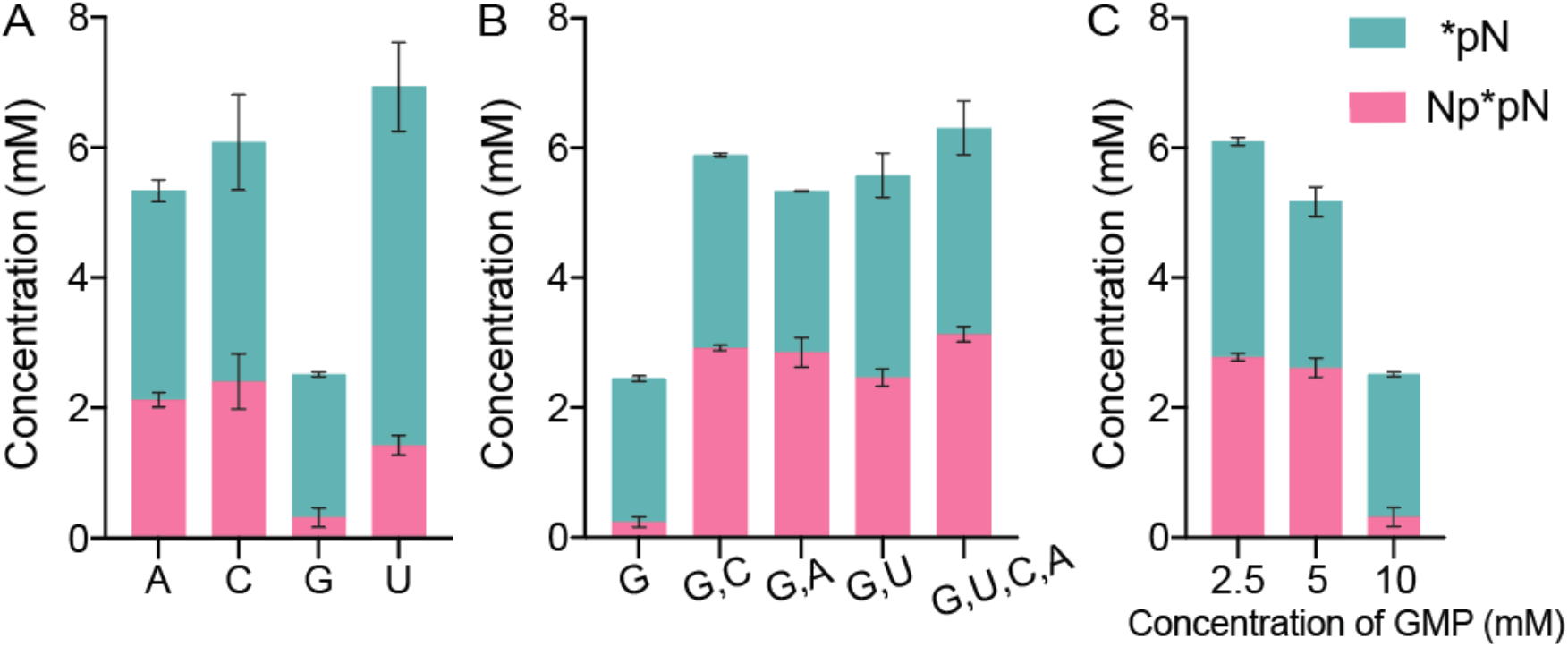
Efficient eutectic ice phase activation of all four canonical nucleotides. (A) Each of the four canonical ribonucleotides could be activated using the same approach as optimized for pA (Figure 1D). However, pG exhibited lower activation compared to the other three. 10 mM pN, 10 mM 2AI, 100 mM 2MBA, 50 mM MeNC, 30 mM MgCl_2_, 50 mM Na^+^-HEPES pH 8, and periodic addition of MeNC in three aliquots of 50 mM each. (B) All four nucleotides can be activated efficiently with minimal side products when incubated simultaneously. Indicated concentrations are sum totals across all the nucleotides. Total [pN] = 10 mM, 10 mM 2AI, 100 mM 2MBA, 50mM MeNC, 30 mM MgCl_2_, 50 mM Na^+^-HEPES pH 8, and periodic addition of MeNC in three aliquots of 50 mM each. (C) GMP activation yields decrease as its concentration is increased, suggesting that the low activation of GMP is partly due to its low solubility in aqueous phase (compare with the other three canonical ribonucleotides in Figure 3A). Total [pN] = 10 mM, 10 mM 2AI, 100 mM 2MBA, 50 mM MeNC, 30 mM MgCl_2_, 50 mM Na^+^-HEPES pH 8, and periodic addition of MeNC in three aliquots of 50 mM each. The complete product breakdown is included in Figure S6. Errors are standard deviations of the mean, n ≥ 2 replicates.

Encouraged by these results, we sought to apply combined ice eutectic phase nucleotide activation and bridged-intermediate formation to template-directed primer extension (Figure 4A). First, we aimed to confirm that nucleotides activated using this approach behave the same way as chemically prepared and purified activated nucleotides. We activated 5 mM each of CMP and GMP under ice eutectic phase conditions (Figure 3B) and then added the resulting mixture to a primer-template duplex in solution phase. Under these conditions, the primer was extended (Figure 4C), just as in the control case with purified chemically prepared activated mononucleotides (Figure 4B). This demonstrates that ice eutectic phase generates activated mononucleotides and bridged dinucleotides, which are then functional in primer extension. Next, we asked whether *in situ* activation and primer extension can occur together in a single reaction mixture during freeze-thaw cycles. In this experiment the primer-template duplex was included along with all other reagents from the start and exposed to cycles of ice eutectic phase activation. We found that primer extension was comparable to that observed with pure activated mononucleotides or with sequential eutectic activation and solution phase primer extension (compare Figure 4E to Figures 4B, 4C/D respectively). We then considered whether this *in situ* pathway could also be combined with our previously-developed bridge-forming activation, where isocyanide and aldehyde are added to activated mononucleotides to generate enhanced levels of bridged dinucleotides from activated mononucleotides in solution and thereby promotes primer extension. We performed *in situ* nucleotide activation together with primer extension as in Figure 4E, but then further treated the mixture with bridge-forming chemistry at room temperature and incubated the mixture for an additional 24 hours. As expected, primer extension was even more efficient, with 52.4±0.3% of the total primer extended to +3 product compared with 38.6±0.7% in the absence of additional bridge-forming activation (Figure 4F). This experiment demonstrates that ice eutectic phase concentration enables nucleotide activation with bridge-forming activation and primer extension, all in one reaction mixture.

**Figure 4.**
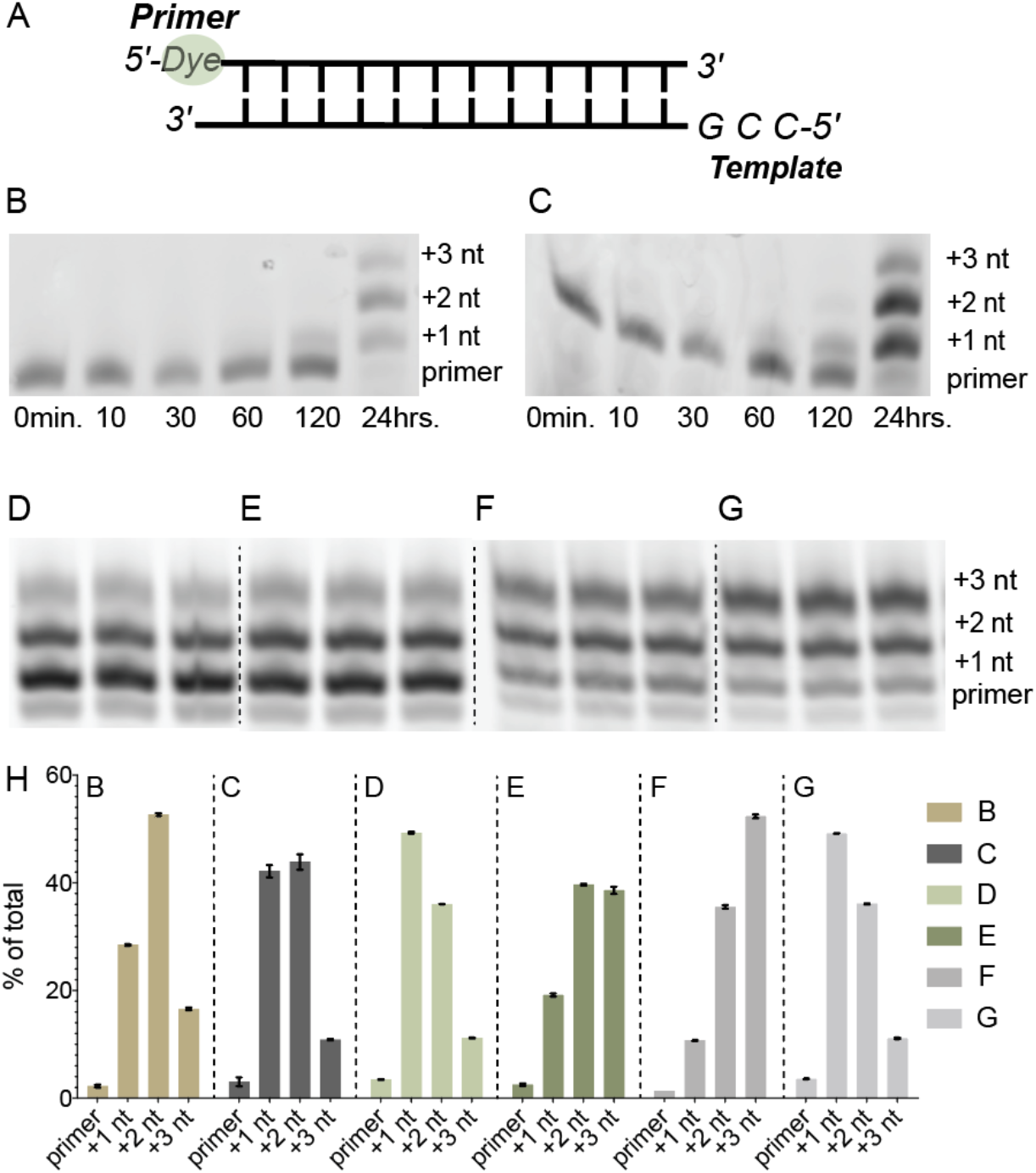
Ice eutectic phase activation enables a complete pathway from unactivated mononucleotides through activation to primer extension. (A) Schematic of primer-template complex. (B) Time course of primer extension using pure activated mononucleotides (5 mM *pG and 5 mM *pC, room temperature). (C) Time course of primer extension reactions using pNs (N = C and G) generated with isocyanide activation chemistry under ice eutectic phase. (D) Primer-template was added to already-activated mononucleotides generated by MeNC-based nucleotide activation under ice eutectic phase conditions, yielding extended primer (as in C). (E) Primer-template duplex was directly treated with pNs (N = C and G) and isocyanide activation chemistry under ice eutectic phase. The products of the reaction were then incubated for an additional 24 hours at room temperature (F) without and (G) with bridge-forming activation. Each set of three lanes in (D-G) represents independent experimental replicates under the indicated condition. (H) Quantification of primer extension products. Positions of primer and +1 to +3-nucleotide (nt) extension products are indicated. Gel image of replicates for (B-C) is included in Figure S9. *Reaction conditions:* (B) 5 mM *pG and 5 mM *pC were added to 1 μM primer, 1.5 μM template, 200 mM Na^+^-HEPES pH 8.0, and 30 mM MgCl_2_. (C and D) 1 μM primer and 1.5 μM template were added to a mixture of 10 mM pN, 10 mM 2AI, 100 mM 2MBA, 30 mM MgCl_2_, 50 mM Na^+^-HEPES pH 8, after four cycles of ice eutectic phase activation (50 mM MeNC each addition). (E) 1 μM primer, 1.5 μM template, 200 mM Na^+^-HEPES pH 8.0, and 30 mM MgCl_2_, 10 mM pNs, 100mM 2MBA, 30 mM MgCl_2_, and periodic addition of MeNC in four aliquots of 50 mM each under ice eutectic phase. The reaction mixture was incubated at room temperature (F) without and (G) with additional 100 mM 2MBA and 200mM MeNC for 24 hours.

The experiments discussed so far used defined templates with only two different nucleobases. To test the limits of our approach with respect to prebiotic plausibility and heterogeneity, we next asked whether the ice eutectic phase activation of all four nucleotides could be combined with bridge-forming chemistry and primer extension to copy random sequence templates. We quantitatively examined the extent and accuracy of this combined process using our recently developed deep-sequencing technique that employs a self-priming RNA hairpin construct with a six-base random-sequence template (see Materials and Methods) (Figure 5A).^15^ The sequencing data reports on both complementary and mismatched nucleotide incorporations resulting from nonenzymatic primer extension. We subjected 1 μM of the RNA sequencing construct along with 10 mM NMPs to four cycles of ice eutectic phase nucleotide activation, and then a 24-hour incubation at room temperature with bridge-forming activation. This reaction extended just 18.4% of the primer by one or more nucleotides, with a 7.9% error frequency (Figure 5B). However these results are closely comparable to primer extension during a 24-hour incubation of the RNA hairpin construct with 10 mM activated nucleotides and bridge-forming chemistry, which results in 19.1% primer extension with 7.3% mismatches (see below for a discussion of why copying in a four-base system is less efficient than in a two-base system), while incubation with 20 mM activated nucleotides in the absence of bridge-forming chemistry results in 11% mismatches.^16^ In both cases we find that complementary products are somewhat G/C-rich at the +1 position, and become further enriched in G and C in downstream positions (Figure 5C). These results are due to G- and especially C-containing bridged dinucleotides being more reactive (Figure 5D). The position-dependent distribution of products, complementary product base preferences, and bridged dinucleotide reactivities are the same as previously observed for primer extension reactions with all four nucleotide reactants.^16^ We conclude that ice eutectic phase activation is compatible with primer extension using all four canonical nucleotides.

**Figure 5.**
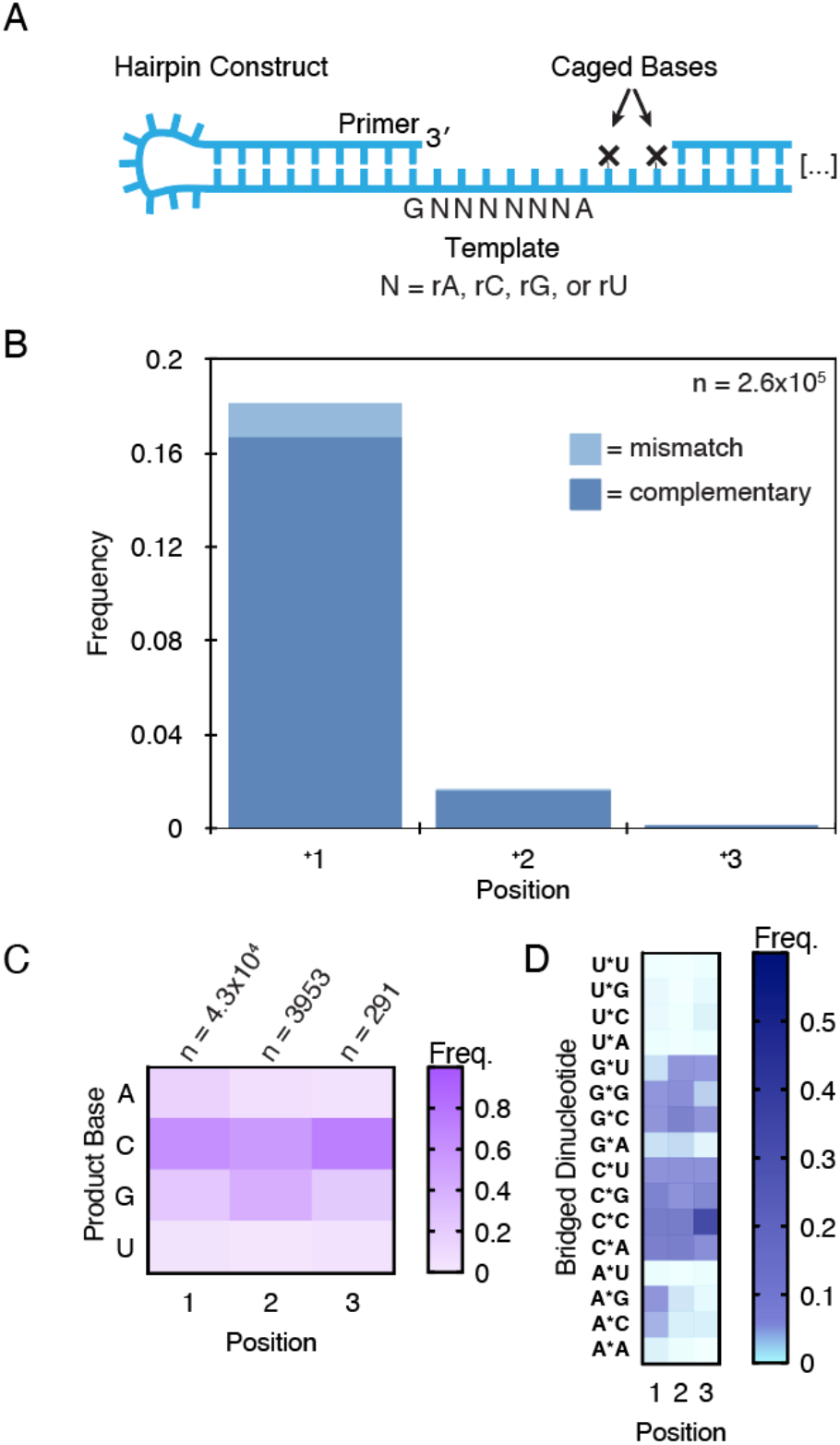
Primer extension on a random sequence template with all four activated nucleotides. (A) A self-priming RNA hairpin construct for deep-sequencing primer extension products on random templates. The caged bases prevent primer extension from encroaching on downstream sequences needed for processing (see Materials and Methods). (Adapted from reference 16) (B) Position-dependent frequencies of mismatched and complementary nucleotide incorporations on a random template as characterized by deep-sequencing. n = unextended hairpins + total nucleotide incorporation events. (C) Position-dependent base frequencies of complementary products. (D) Position-dependent frequencies of bridged dinucleotides that participated in generating complementary products.

To better discern the contribution of ice eutectic phase incubations to the generation of primer extension products, we also tested activating the nucleotides first and then subsequently incubating them with the sequencing hairpin for 24 hours. In the absence of any additional bridge-forming chemistry, the product yields were lower than above (12%) and the error frequency higher (9%) (Figure S7A). If the hairpin was included during the ice eutectic phase activation but the bridge-forming chemistry omitted during the 24-hour room temperature incubation, the yield was slightly lower (14%) and the error frequency slightly higher (8.4%) compared to the experiment above in which bridge-forming chemistry was included (Figure S7B). Finally, if the nucleotides were activated first and then added to the hairpin with bridge-forming chemistry, the yield (17%) and fidelity (7% mismatched) were comparable to the experiment above (Figure S7C). Taken together, these experiments indicate that including the primer-template during the ice eutectic phase incubations with methyl isocyanide and aldehyde does not significantly affect the final products of primer extension, and that bridge-forming chemistry consistently increases yields and improves fidelity.

We have previously found that the error frequency of primer extension products is determined by the ratio of bridged dinucleotides to activated mononucleotides, with a higher ratio predictive of lower error frequencies.^16^ Compared to error frequencies found without any chemistry to drive bridged dinucleotide formation, the values reported here are all relatively low. This is consistent with the high ratio of bridged dinucleotides to activated mononucleotides measured by NMR after cycles of eutectic activation (Figure S8A). Finally, the pattern of mismatches is comparable between experiments whether the hairpin was included during the ice eutectic phase cycles or added subsequently (Figure S8B-C). We conclude that neither the ice eutectic phase conditions nor the prolonged exposure to the activating reagents increase error frequencies or otherwise impose any significant detriment on primer extension.

## Discussion

The plausibility of any scenario for the origin of life on the early Earth is greatly enhanced if different steps or stages in the overall process can occur together under mutually compatible physical and chemical conditions. There are many examples of potentially prebiotic chemical reactions that at first appeared to be incompatible but were subsequently reconciled. For example, it used to be thought that the oxygenous chemistry leading from formaldehyde to sugars such as ribose, and the nitrogenous chemistry leading from cyanide to the nucleobases were strictly incompatible due to the formation of unreactive glycolonitrile and therefore had to occur in separate environments. However, the cyanosulfidic reaction network studied by Sutherland and colleagues has shown that this is not the case, and that it is actually advantageous for these steps to occur together because the facile reduction of glycolonitrile makes it an excellent starting point for many synthetic reactions.^17-19^

Resolving apparent incompatibilities between distinct aspects of prebiotic chemistry, and especially between sequential chemical reactions, is a significant challenge in origin of life studies, where the goal is to elucidate robust and realistic pathways leading from simple chemical feedstocks to replicating and evolving protocells. In the present work, we have focused on one such incompatibility, namely the very high concentrations of the activating moiety 2-aminoimidazole (2AI) thought to be required for nucleotide activation, and the fact that such high concentrations of 2AI prevent the accumulation of the bridged dinucleotide required for efficient template directed RNA copying. We have found that ice eutectic phase incubation enables nucleotide activation with equimolar concentrations of 2AI and nucleotides, thus allowing bridged dinucleotides to accumulate under prebiotically plausible scenarios. Furthermore, cycles of MeNC addition and eutectic freezing drive both efficient activation and bridged dinucleotide formation, directly enabling efficient primer extension. Thus, an environment subject to repeated cycles of freezing and thawing enables the operation of a potentially prebiotic pathway that begins with ribonucleotide 5′-monophosphates, generates 2AI activated nucleotides and then the essential bridged dinucleotides, and finally yields primer extension products, all in one mutually compatible system.

Our deep-sequencing results show that the multi-step pathway from activation to primer extension is compatible with the presence of all four canonical nucleotides, but the very low overall yield of primer extension products on templates containing all four canonical nucleotides strongly suggests that some aspect of the copying chemistry is limiting. The low yield of primer extension products with all four canonical activated mononucleotides (as opposed to just G and C) and the corresponding bridged dinucleotides is a known problem.^16^ For example, formation of ten possible bridged dinucleotides from four canonical nucleotides results in lower relative concentrations of each individual bridged dinucleotide. In addition, overlapping binding sites on the template may decrease the fraction bridged dinucleotides correctly bound next to primer. However, previous experiments have shown that activated helper oligonucleotides can enable more extensive template copying,^3^ while noncanonical but prebiotically plausible nucleotides such as 2-thio-U greatly enhance the copying of A and U residues in the template strand.^20^ We are currently investigating whether the presence of helper oligonucleotides, or replacing one or more of the canonical nucleotides with prebiotically plausible alternative nucleotides might lead to more efficient template copying.

Our observation that the initial activation of mononucleotides can occur under ice eutectic phase conditions suggests a potential role for icy early Earth environments in nonenzymatic RNA copying. The formation and thawing of ice is a common phenomenon on Earth’s surface, which can take place during day-night temperature cycles, seasonal temperature changes, or random weather-based changes.^21-22^ Our results are consistent with a potentially wide role for freeze thaw cycles in promoting processes from prebiotic synthesis^10^ to RNA copying, and may even have astrobiological implications.^23-24^ We observed that the efficiency of nucleotide activation is robust with respect to changes in the length of freeze-thaw cycles and the frequency of MeNC addition. MeNC is thought to have been released from prebiotic feedstocks following exposure to UV irradiation from the Sun.^5^ The relative independence of activation efficiency from the frequency of MeNC delivery suggests that the required temperature cycling conditions are not stringently constrained, and therefore any corresponding early Earth environment need not be highly contrived. Ice eutectic-based nucleotide activation may not even have been required continuously: once the initial pool of activated mononucleotides has been generated, the application of solution-phase bridge-forming activation could slowly re-activate hydrolyzed nucleotides as well as regenerate bridged dinucleotides in solution. Additional experiments will be needed to determine how long this system can replenish hydrolyzed activated nucleotides without the need for additional freezing after the initial activation.

The key ingredient in the chemical nucleotide activation procedure that we used is MeNC, so it is important to consider whether MeNC could be generated in an environment compatible with our pathway. The strong UV absorption by nucleotides might leave insufficient UV light for MeNC generation. However, nucleotides mostly absorb light between 240 nm and 280 nm, whereas the generation of MeNC proceeds with longer wavelength irradiation.^25^ Specifically, two steps in the plausible synthesis of MeNC exploit the use of UV light: the production of nitroprusside has been achieved with 365 nm or 254 nm light and isocyanide ligand exchange also occurs during exposure to 365 nm light,^5^ consistent with the broad band irradiation from the young Sun.^26^ The formation of MeNC using longer wavelength UV irradiation suggests that the generation of MeNC is compatible with the presence of nucleotides. Another consideration is that an important ingredient in MeNC synthesis is methylamine, which could have been delivered to the primordial Earth by comets, or by reduction of HCN with hypophosphite and sponge nickel.^5, 27^ On the basis of Ugi-type reactions,^28^ a primary amine such as methylamine will readily react with an aldehyde, forming an iminium ion, however, this may not affect the nucleotide activation chemistry because MeNC will still add reversibly to the iminium ion to produce the nitrilium ion required for nucleotide activation.^12^ In addition, 2-hydroxy-N-methylpropanamide, the side product of nucleotide activation, will hydrolyze to regenerate methylamine, which can then take part in another round of MeNC synthesis. ^5^ The prebiotic plausibility of the remaining chemical ingredients has been previously discussed,^17, 19, 29^ but additional experiments will be needed to reconstitute in situ MeNC generation with simultaneous nucleotide activation.

Continuous re-activation chemistry may eventually enable multiple cycles of nonenzymatic RNA replication. Maintaining template-directed polymerization activity for long periods requires the population of mononucleotides to be in a highly activated state (a high ratio of bridged dinucleotides to 5′-phosphorimidazolides to 5′-phosphates) in the face of spontaneous hydrolysis. Further work will be necessary to determine the required frequency of ice eutectic activation to enable continuous template copying and ultimately RNA replication. Importantly, the presence of the primer/template complex during the ice eutectic phase incubation with MeNC and aldehyde does not result in higher frequencies of mismatches, suggesting that these reagents do not generate mutagenic lesions and that the low temperature does not favor misincorporation events. It is possible that although the ice eutectic phase concentration favors activation chemistry, the low temperature may inhibit the actual primer extension reaction. Indeed, primer extension during the freeze-thaw phase of our experiments may be occurring primarily during the relatively short but non-negligible thaw intervals. The inclusion of bridge-forming chemistry during a subsequent liquid phase incubation has a more pronounced effect on mismatch frequencies, presumably because increasing the ratio of bridged dinucleotides to activated mononucleotide increases the fidelity of templated copying, as previously demonstrated.^7^ Nucleotide activation during cycles of partial drying and rehydration may also be relevant.^30^

## Materials and Methods

### 1.1 General information

#### Materials

Reagents and solvents were obtained at the highest available purity from Acros Organics, Alfa Aesar, Fisher Scientific, Sigma-Aldrich, ThermoFisher Scientific, or Tokyo Chemical Industry Co., and were used without further purification unless noted below. Nucleoside-5′-monophosphates, free acid, were purchased from Santa Cruz Biotechnology. 2-aminoimidazole hydrochloride and 2,2′-dipyridyldisulfide were purchased from Combi Blocks. RNA oligonucleotides for standard primer extension experiments were purchased from Integrated DNA Technologies. RNA oligonucleotides used for deep-sequencing experiments were prepared as described in Section 1.5. The synthesis and storage of methyl isocyanide was as previously described.^7^ All reactions were carried out in DNase/RNase-free distilled water.

#### NMR spectroscopy

^1^H and ^31^P-NMR spectra were obtained using a Varian INOVA NMR spectrometer operating at 400 MHz and 161 MHz respectively. Samples in H_2_O/D_2_O mixtures were analyzed using Wet1D suppression to collect ^1^H-NMR data. Chemical shifts (δ) are shown in ppm. Coupling constants (J) are given in Hertz (Hz) and the notations s, d, t, and m represent singlet, doublet, triplet, and multiplet multiplicities respectively.

#### pH measurements

pH values were determined by a micro pH probe (Orion 9863BN) equipped with a needle tip and a SevenCompact meter (Mettler Toledo S220).

#### Data analysis

All NMR spectra were analyzed using MestReNova (version 12.0.3). The yields of conversion were determined by the relative integration of the signals in the ^1^H or ^31^P NMR spectra.

### 1.2 Preparation, storage, and concentration determination of stock solutions

Stock solutions of nucleoside-5′-monophosphate disodium salt and 2-aminoimidazole hydrochloride were prepared in DNase and RNase-free distilled water. After adjusting the pH to the reported values with NaOH/HCl, the stock solution was filtered with 0.22-micron syringe filters (Millipore Sigma). Each stock solution was then aliquoted and kept at -20 °C until further use. The concentrations of the nucleoside-5′-monophosphate solutions were determined by analysis of serial dilutions on a UV spectrophotometer. The absolute concentrations of other stock solutions were determined by comparing the integrals of ^1^H-NMR peaks of interest to the calibrant, trimethyl phosphate, by NMR spectroscopy.

### 1.3 Sample preparation for ice eutectic phase experiments

Reaction mixtures were first flash-frozen through by immersing the plastic tubes in liquid nitrogen, followed by incubation at -13 °C in a standard laboratory freezer unit for the indicated time. ^9, 13, 31^

### 1.4 Primer extension reactions and product analysis

#### Control Primer Extension Reactions

The primer-template duplex was first annealed in a 40 μl solution containing 2 μM thiol-modified primer (/5ThioMC6-D/AGUGAGUAACUC, IDT), 3.1 μM template (CCGGAGUUACUCACU, IDT), 125 mM Na^+^-HEPES (pH 8.0), and 62.5 mM MgCl_2_ by heating at 95 °C for 3 min followed by cooling to 23 °C at a rate of 0.1 °C /s. The reaction was initiated by the addition of 10 μl 100 mM activated mononucleotides (pH ∼9.6).

#### Sample Purification and preparation (adapted from reference ^7^)

10 μl aliquots were removed at given time points and desalted using a ZYMO Oligo Clean & Concentrator spin column (ZYMO Research). The isolated material was resuspended in 30 μl 100 mM Na^+^-HEPES (pH 7.50) and disulfide bonds were reduced using a 10-fold molar excess of tris-(2-carboxyethyl) phosphine hydrochloride (TCEP). Alexa 488 C5 maleimide dissolved in anhydrous dimethyl sulfoxide (DMSO) at a concentration of 1 mM was added to the primer-template duplex dropwise, and the coupling reaction was allowed to proceed in the dark at room temperature for 2 hours. The labeled primer-template duplex was separated from free dye using ZYMO DNA Clean & Concentrator spin columns, resuspended in 5 μl of 100 mM Na^+^-HEPES buffer, and mixed with 30 μl of quenching buffer containing 8.3 M urea, 1.3x Tris/Borate/EDTA (TBE) buffer (pH 8.0), 0.005% Bromphenol Blue, 0.04% Orange G, and 75 μM RNA complementary to the template. To denature the labeled product from the template prior to polyacrylamide gel analysis, and sequester the template with its complement (added in excess as a competitor), the sample was heated to 95 °C for 3 min and cooled to 23 °C at 0.1 °C/s.

#### Analysis of the polymerization products

20% polyacrylamide gels were prepared using the SequaGel–UreaGel system (National Diagnostics, Atlanta, GA). The gels were allowed to pre-run at a constant power of 20 W for 40−45 mins. 15 μl aliquots of samples were separated by 20% (19:1) denaturing PAGE at a constant power of 5 W for 10 min. and 25 W for 1.5 hrs. Gels were imaged on a Typhoon 9410 scanner (GE Healthcare, Little Chalfont, Buckinghamshire, UK) and quantified using the accompanying ImageQuant TL software. All reactions were performed in triplicate or greater.

### 1.5 RNA sample preparation for deep-sequencing (adapted from reference ^16^)

Reagents, reaction conditions, the NERPE-Seq protocol, and the data analysis code package were as previously described.^15^ Na^+^-HEPES buffer was prepared from the free acid (Sigma-Aldrich), adjusted to pH 8 with NaOH, and filtered. Enzymes were purchased from New England BioLabs (NEB). Incubations at a specified temperature were performed in a Bio-Rad T100 thermal cycler, excepting reactions with methyl isocyanide which were carried out at room temperature (∼23 °C).

1 μM of the sequencing hairpin construct (GUUCAGAGUUCUACAGUCCGACGAUCdT(-NPOM)CdT(-NPOM)ANNNNNNGCAUGCGACUAAACGUCGCAUGC, dT(-NPOM) = NPOM-caged deoxyT, ^32^ N = rA, rU, rC or rG) and 1.2 μM 5’
s Handle Block (GUCGGACUGUAGAACUCUGAA-dideoxyC, IDT) were annealed by incubation at 95°C for 3 minutes, and cooling to 23 °C at 0.2 °C/s. The activated ribonucleotides, chemical reactants, and 30 mM MgCl_2_ were added to initiate the reaction. The mixture was briefly vortexed and incubated at ∼23 °C or subjected to ice eutectic phase freezing and thawing, as indicated. A 30 μl aliquot of a given reaction was quenched by the addition of 20 μl water followed by desalting in a MicroSpin G-25 spin column (GE Healthcare). The NPOM (caged) bases were uncaged by a 45-minute exposure under a 385 nm UV lamp (∼3 cm distance from the tops of the tubes, Spectroline ENF-240C, Spectronics). Samples were PAGE-purified (ZR small-RNA PAGE Recovery Kit, Zymo Research). The RT Handle (template for the reverse transcription primer, App-AGATCGGAAGAGCACACGTCT-dideoxyC, App = riboA 5′-adenylation, IDT) was ligated with T4 RNA Ligase 2, truncated KQ. The mixture was treated with Proteinase K, phenol-chloroform extracted, and concentrated with an Oligo Clean & Concentrator spin column (Zymo Research).

The primer for reverse transcription (AGACGTGTGCTCTTCCGATCT, IDT) was annealed to the purified sample and the RNA reverse transcribed with ProtoScript II. The mixture was purified with an Oligo Clean & Concentrator spin column, and the eluted cDNA stock concentration measured by spectrophotometry. 0.1 μg cDNA was added to a 50 μl Q5 Hot Start High-Fidelity DNA Polymerase PCR reaction with 0.2 μM each of NEBNext SR Primer for Illumina and NEBNext Index Primer for Illumina (NEB) and run for 6 cycles with a 15 s 62 °C extension step. PCR product was purified by preparative agarose gel electrophoresis (Quantum Prep Freeze ‘N Squeeze spin column, Bio-Rad) and magnetic beads (Agencourt AMPure XP, 1.7:1 volume ratio of magnetic bead suspension to sample volume). Samples were validated and concentrations measured by TapeStation (Agilent). Paired-end sequencing by MiSeq (Illumina, MiSeq Reagent Kit v3, 150 cycle) produced ∼20 million reads with ∼95% passing the instrument’s quality filter. Previous work has evaluated the effects of 2′-5′ linkages on the sequencing protocol and found that they do not measurably alter the reported sequencing data. The experimentally established error associated with each reported base identity is 0.062 ± 0.13%.^15^

Sequencing data analysis was performed with the NERPE-Seq custom code package written in MATLAB (MathWorks) as previously described. ^15^ Briefly, the code filters raw data by using read quality scores, and by checking that reads agree with defined-sequence regions of the hairpin construct and that forward and reverse reads (which overlap) agree with each other. Template and product sequence pairs are extracted and characterized. The template sequence of the construct does not contain precisely equal fractions of rA, rC, rG and rU, ^15^ so the template from a control experiment in which no activated nucleotides were added was used to generate normalization factors. All presented data are normalized—showing results as they would appear if the template were perfectly random with equal base ratios. The numerical data used to generate heat maps is included below.

## Data Availability Statement

The NERPE-Seq analysis code is available in the GitHub repository: https://github.com/CarrCE/NERPE-Seq.

Raw sequencing data and NERPE-Seq analysis output data are available at OSF.io: https://osf.io/7sxgz/?view_only=4f665338d1474855940b13fba1dfb53d

Raw NMR data is available at OSF.io: https://osf.io/sjtwu/?view_only=c38e3a8bced14c2880fb5323d9f12891

## Acknowledgments

J.W.S. is an Investigator of the Howard Hughes Medical Institute. This work was supported in part by grants from the Simons Foundation (290363) and the NSF (CHE-1607034) to J.W.S and from NASA (80NSSC19K1028) to C.E.C. The authors thank members of the Szostak laboratory for helpful discussions, and comments on the manuscript.

