## Supplementary Information for "Freeze-thaw Cycles Enable a Prebiotically Plausible and Continuous Pathway from Nucleotide Activation to Nonenzymatic RNA Copying"

* Jack W. Szostak

**This PDF file includes:**

Figures S1 to S9
Tables S1 to S4

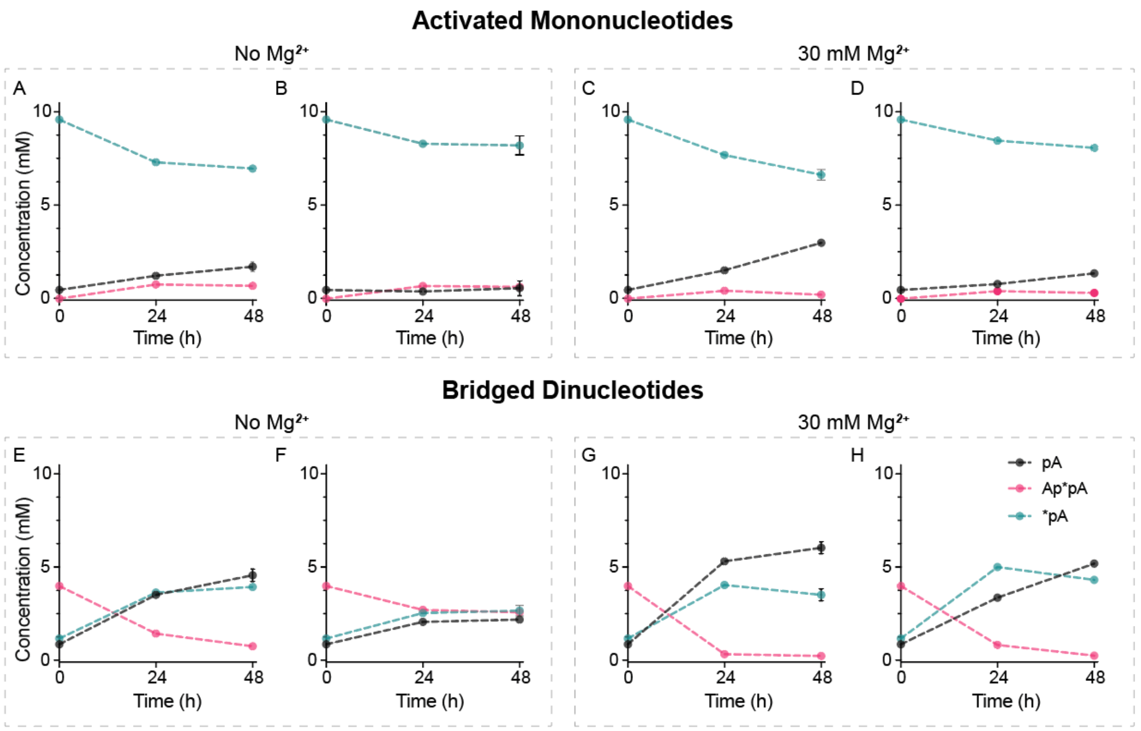


**Figure S1.** Activated nucleotides (*pA) are less susceptible to hydrolysis under ice eutectic conditions at -13 °C compared to room temperature. In the absence of $Mg^{2+}$, more *pA hydrolyzed to pA at (A) room temperature compared to (B) ice eutectic phase. A similar level of bridged dinucleotides is formed under either condition. With 30 mM $Mg^{2+}$, more *pA hydrolyzed to pA at (C) room temperature compared to (D) ice eutectic phase. In addition, the presence of $Mg^{2+}$ accelerated the hydrolysis either under room temperature (compare (A) and (C)) or ice eutectic phase (compare (B) and (D)). In the absence of $Mg^{2+}$, there is almost no Ap*pA left under (E) room temperature while over 50% of Ap*pA remained under (F) ice eutectic phase. The presence of 30 mM $Mg^{2+}$ significantly increased the hydrolysis rate of Ap*pA under either (G) room temperature or (H) ice eutectic phase. However, importantly, at the 24-hr timepoint, less Ap*pA has been hydrolyzed under the ice eutectic phase (H) compared to (G) room temperature. Note that the effective concentration of $Mg^{2+}$ in the ice eutectic phase might be much higher than 30 mM. *Reaction conditions*: 10 mM *pA, 200 mM $Na^+$-HEPES at pH 8 under (A) room temperature and (B) ice eutectic phase at -13 °C. 10 mM *pA, 30 mM $Mg^{2+}$, 200 mM $Na^+$-HEPES at pH 8 under (C) room temperature and (D) ice eutectic phase. 10 mM Ap*pA, 200 mM $Na^+$-HEPES at pH 8 under (E) room temperature and (F) ice eutectic phase. 10 mM Ap*pA, 30 mM $Mg^{2+}$, 200 mM $Na^+$-HEPES at pH 8 under (G) room temperature and (H) ice eutectic phase.
Errors are standard deviations of the mean, n ≥ 2 replicates.

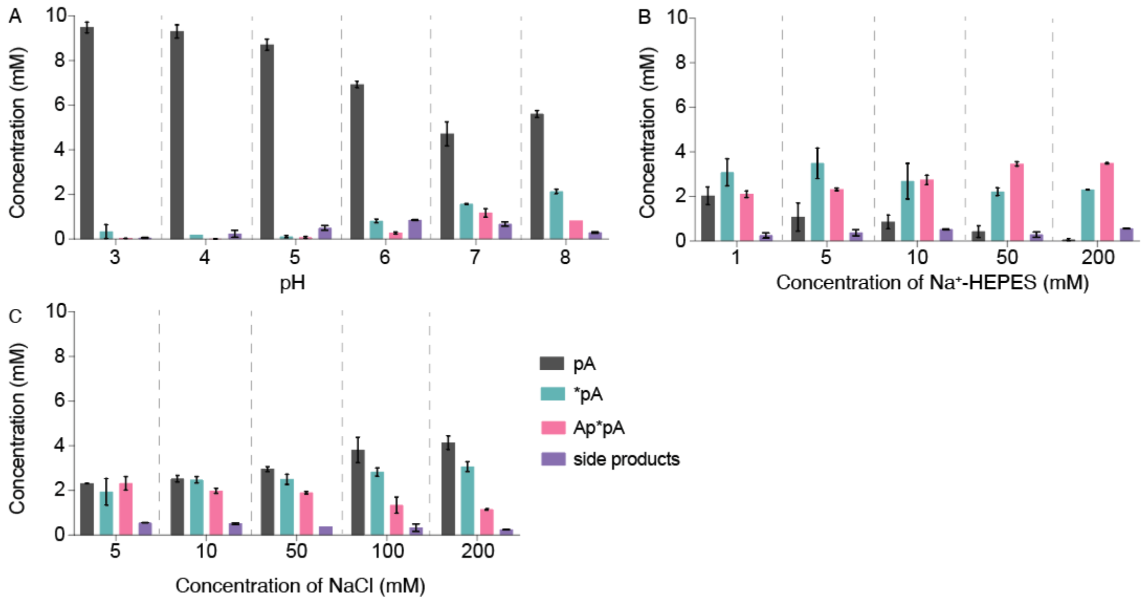


**Figure S2.** The importance of pH effects under ice eutectic phase. (A) Eutectic activation was performed across a range of initial pH values (3 to 8). The solution was brought to the indicated pH by addition of concentrated HCl or NaOH. No buffering agent was used. The optimal initial pH is 7. The low yields of activation compared to the case in which $Na^+$-HEPES buffer is used (compare to (B)) suggests that the initial adjustment of the pH is not sufficient to regulate the pH during the incubation under ice eutectic phase. (B) Increasing $Na^+$-HEPES (up to 200 mM) increased the activation yield. Note that significant activation is observed with only 1 mM $Na^+$-HEPES, much less than the concentrations of MeNC and 2MBA, suggesting that $Na^+$-HEPES is not actively participating in the isocyanide activation. (Note that the imidazolide moiety of 2AI can provide buffering, however, 2AI is consumed during the reaction over time.) (C) To confirm that the increase in reaction yield in the presence of $Na^+$-HEPES is not due to a simple salt effect, we titrated NaCl into the reaction. Increasing the concentration of NaCl leads to an opposite trend compared to that observed when increasing $Na^+$-HEPES, suggesting that simple ionic effects are not responsible for facilitating the reaction. We note that chloride can strongly affect the pH in the eutectic phase since it is retained in the ice, drawing $H^+$ into the ice for charge balance, making the liquid phase more alkaline.[1] *Reaction conditions*: all experiments were performed using the same approach with four cycles as optimized for pA and shown in Figure 1D. (A) 10 mM pA, 10 mM 2AI, 100 mM 2MBA, 30 mM $MgCl_2$, 50 mM MeNC plus periodic addition of the MeNC under ice eutectic phase. (B) 10 mM pA, 10 mM 2AI, 100 mM 2MBA, 30 mM $MgCl_2$, 50 mM MeNC plus periodic addition of the MeNC, with indicated $Na^+$-HEPES concentration at pH 8 under ice eutectic phase. (C) 10 mM pA, 10 mM 2AI, 100 mM 2MBA, 30 mM $MgCl_2$, 50 mM MeNC plus periodic addition of the MeNC, 5 mM $Na^+$-HEPES pH 8, and indicated amount of NaCl under ice eutectic phase. Errors are standard deviations of the mean, $n \geq 2$ replicates.

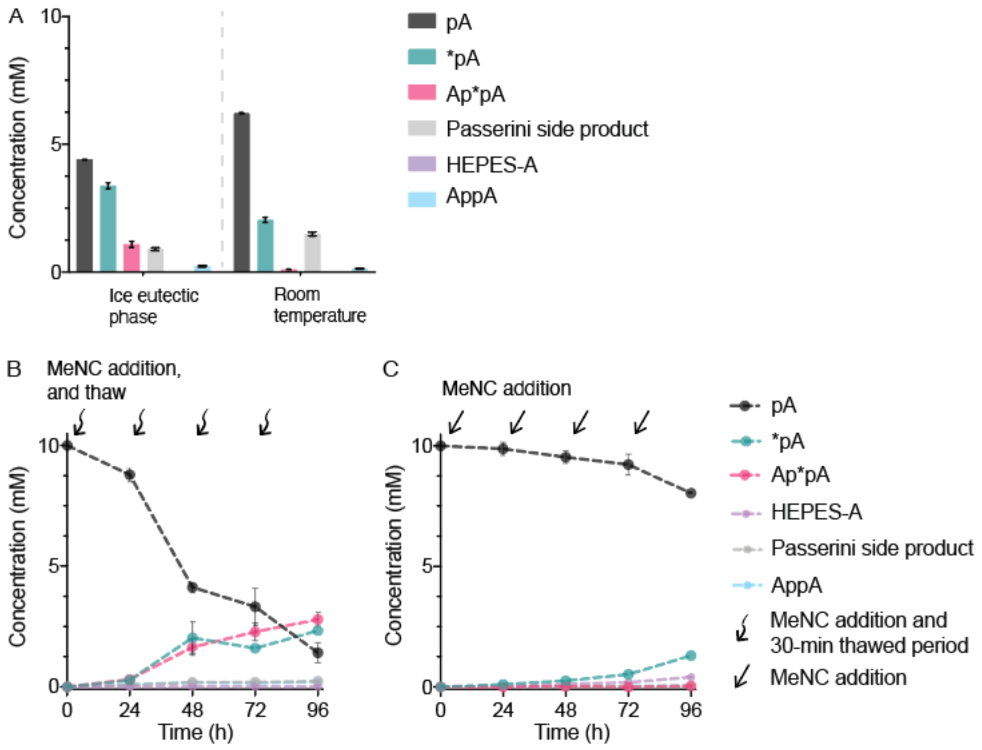


**Figure S3.** Ice eutectic phase significantly enhances the yield of activated nucleotides (*pN) and promotes formation of bridged dinucleotides (Np*pN). The same data as in Figure 1 C-E, but with a complete breakdown of the identities of the side products. (A) Effect of ice eutectic phase on nucleotide activation. Nucleotide activation at ice eutectic phase (left) and room temperature (right) with one round of MeNC addition and equimolar concentrations of initially unactivated nucleotides and 2AI. (B) Cycles of MeNC addition and eutectic freezing drive significant nucleotide activation. Controlled addition of MeNC to the thawed solution enhances the yield of activated nucleotides (*pA). The curvy arrows indicate the addition of MeNC after thawing of the reaction mixture. Note that the reaction mixture is frozen as eutectic ice for the entire reaction course except during the brief thawing steps (~30 minutes at room temperature for the mixture to thaw completely). (C) Activation is inefficient in the absence of ice eutectic freezing, at room temperature. The straight arrows indicate the addition of MeNC at room temperature. *Reaction conditions*: 10 mM pA, 10 mM 2AI, 100 mM 2MBA, 30 mM $MgCl_2$, 50 mM $Na^+$-HEPES pH 8, plus (A) one addition of 200 mM MeNC under ice eutectic phase or room temperature, or (B) periodic addition of the MeNC beyond the initial 50 mM in three aliquots of 50 mM each under ice eutectic phase at -13 °C or (C) at room temperature. Errors are standard deviations of the mean, n ≥ 2 replicates.

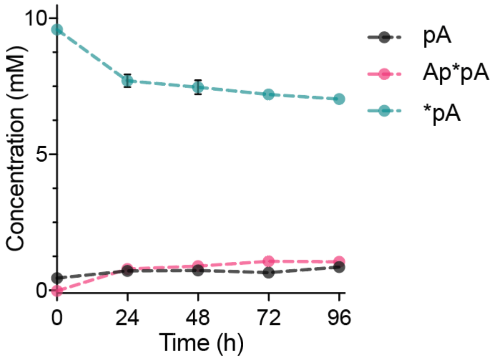


**Figure S4.** Spontaneous bridged dinucleotide formation from pure activated nucleotide (*pA) under ice eutectic phase. The concentration of bridged dinucleotides (Ap*pA) slowly increased over time as 10 mM of *pA was incubated under ice eutectic phase in the absence of isocyanide-driven activation chemistry (Figure S3B). The reaction mixture was under ice eutectic phase conditions except during the four brief thawing steps to make measurements. *Reaction conditions:* 10 mM *pA, 50 mM $Na^+$-HEPES pH 8, 30 mM $MgCl_2$, under ice eutectic phase. Errors are standard deviations of the mean, n = 2 replicates.

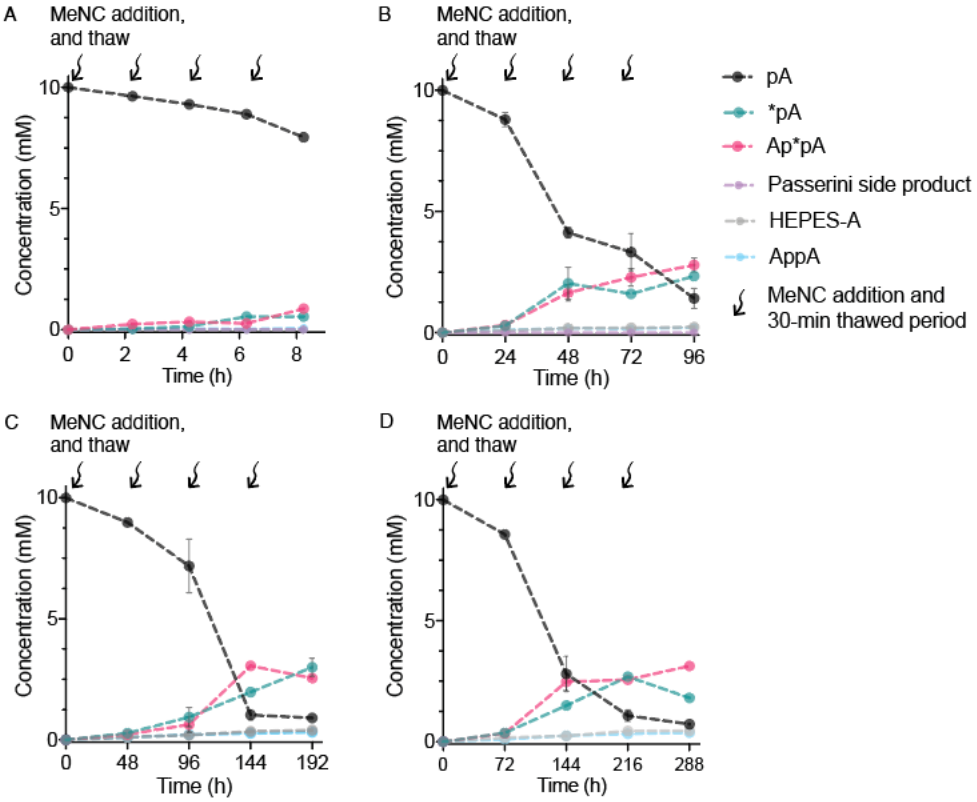


**Figure S5.** The frequency of MeNC delivery at long time intervals does not affect the overall yield of activated products. The same data as in Figure 2, but with a complete breakdown of the identities of the side products. (A) Yields of activated species are low with time intervals on the order of hours in between MeNC additions. (B-D) However, lengthening of the time interval beyond 24 hours does not significantly affect the final yield. The complete product breakdown is included in Figure S5. *Reaction conditions*: 10 mM pA, 10 mM 2AI, 100 mM 2MBA, 50 mM MeNC, 30 mM $MgCl_2$, 50 mM $Na^+$-HEPES pH 8, and subsequent periodic addition of MeNC in three aliquots of 50 mM. Errors are standard deviations of the mean, n ≥ 2 replicates.

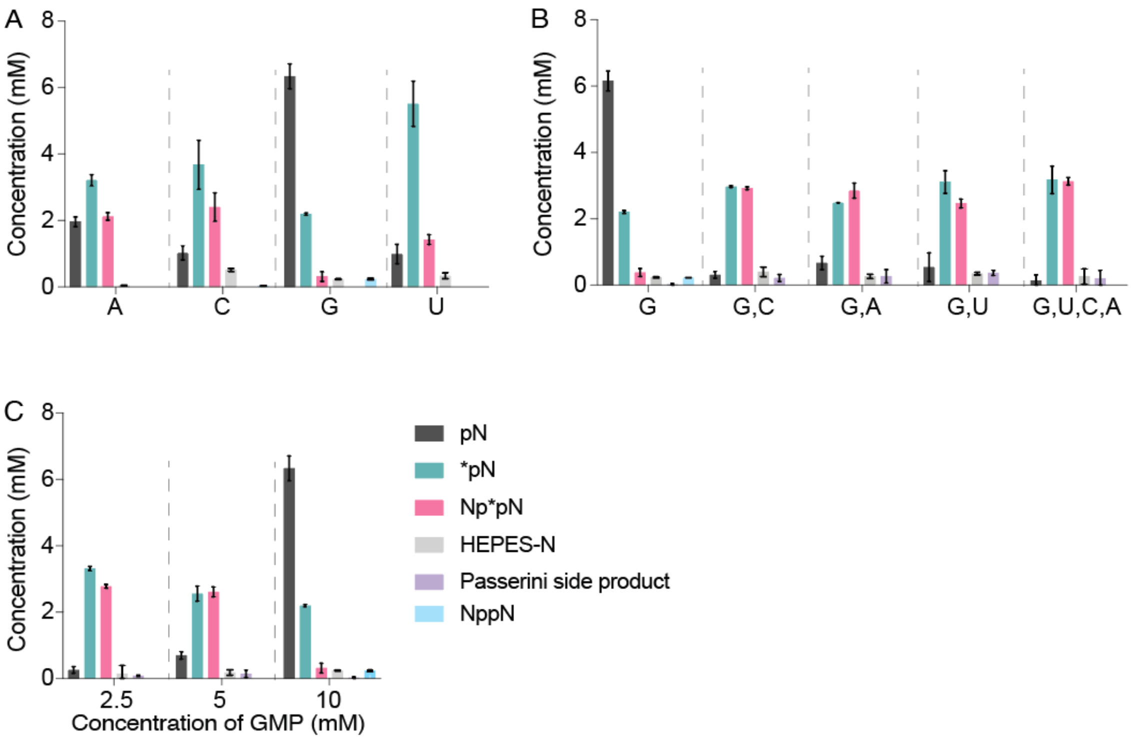


**Figure S6.** Efficient eutectic ice phase activation of all four canonical nucleotides. The same data as in Figure 3, but with a complete breakdown of the identities of the side products. (A) Each of the four canonical ribonucleotides could be activated using the same approach as optimized for pA in Figure 1D. However, pG exhibited lower activation compared to the other three. 10 mM pN, 10 mM 2AI, 100 mM 2MBA, 50 mM MeNC, 30 mM $MgCl_2$, 50 mM $Na^+$-HEPES pH 8, and periodic addition of MeNC in three aliquots of 50 mM each. (B) All four nucleotides can be activated to almost full yield with minimal side products when incubated simultaneously. Indicated concentrations are sum totals across all the nucleotides. Total [pN] = 10 mM, 10 mM 2AI, 100 mM 2MBA, 50mM MeNC, 30 mM $MgCl_2$, 50 mM $Na^+$-HEPES pH 8, and periodic addition of MeNC in three aliquots of 50 mM each. (C) GMP activation yields decrease as GMP concentration is increased, suggesting that its low activation is partly due to its low solubility in aqueous phase. Total [pN] = 10 mM, 10 mM 2AI, 100 mM 2MBA, 50 mM MeNC, 30 mM $MgCl_2$, 50 mM $Na^+$-HEPES pH 8, and periodic addition of MeNC in three aliquots of 50mM each. Errors are standard deviations of the mean, $n \geq 2$ replicates.

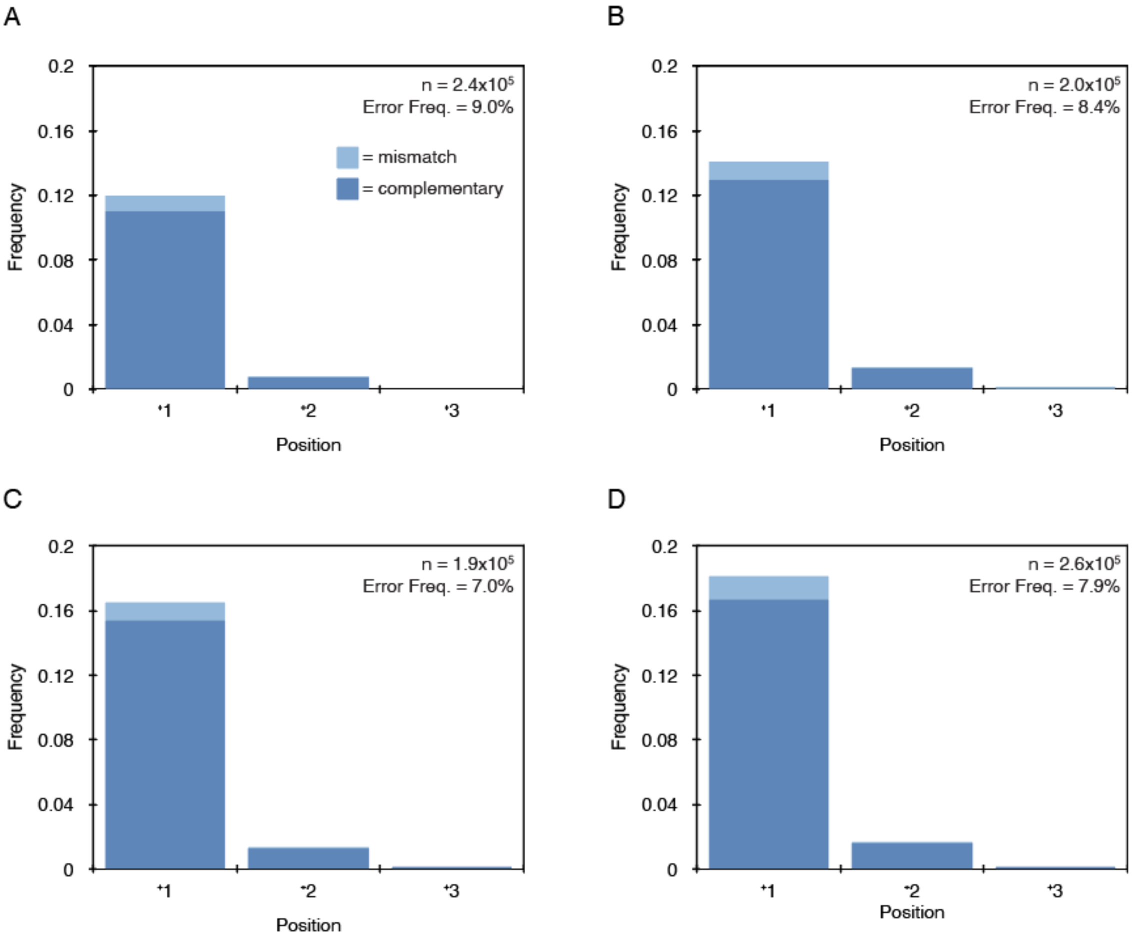


**Figure S7.** Eutectic phase activation is compatible with primer extension using all four nucleotides. (A-D) Position-dependent frequencies of mismatched and complementary nucleotide incorporations on a random template as characterized by deep-sequencing. n = unextended hairpins + total nucleotide incorporation events. Error freq. = percentage of mismatched incorporations relative to all incorporations. (A) Activated nucleotides generated by four rounds of eutectic activation were added to the sequencing hairpin and incubated for 24 hours. (B) The sequencing hairpin was included during the four rounds of eutectic activation, and then incubated for 24 hours. (C) Activated nucleotides generated by four rounds of eutectic activation were added to the sequencing hairpin, and then incubated for 24 hours with bridge-forming chemistry. (D) The sequencing hairpin was included during the four rounds of eutectic activation, and then incubated for 24 hours with bridge forming chemistry. Same as Figure 5B.

Inclusion of the hairpin during the eutectic activation slightly increases yields and does not have a significant negative effect on fidelity (compare (A) and (C), hairpin not included, with (B) and (D), hairpin included). Addition of bridge-forming chemistry increased yields (compare (A) and (B) to (C) and (D).

*Reaction conditions:* 2.5 mM each rNMP (10 mM total), 30 mM $MgCl_2$, 50 mM $Na^+$-HEPES pH 8, 100 mM 2MBA, 50 mM MeNC, and 1 µM sequencing hairpin (where indicated) were subjected to four 24 hour rounds of eutectic phase activation with the addition of 50 mM MeNC at each thaw. The subsequent 24-hour incubation was at 23°C. Bridge-forming chemistry, where indicated, was the addition of 100 mM 2MBA and 200 mM MeNC.

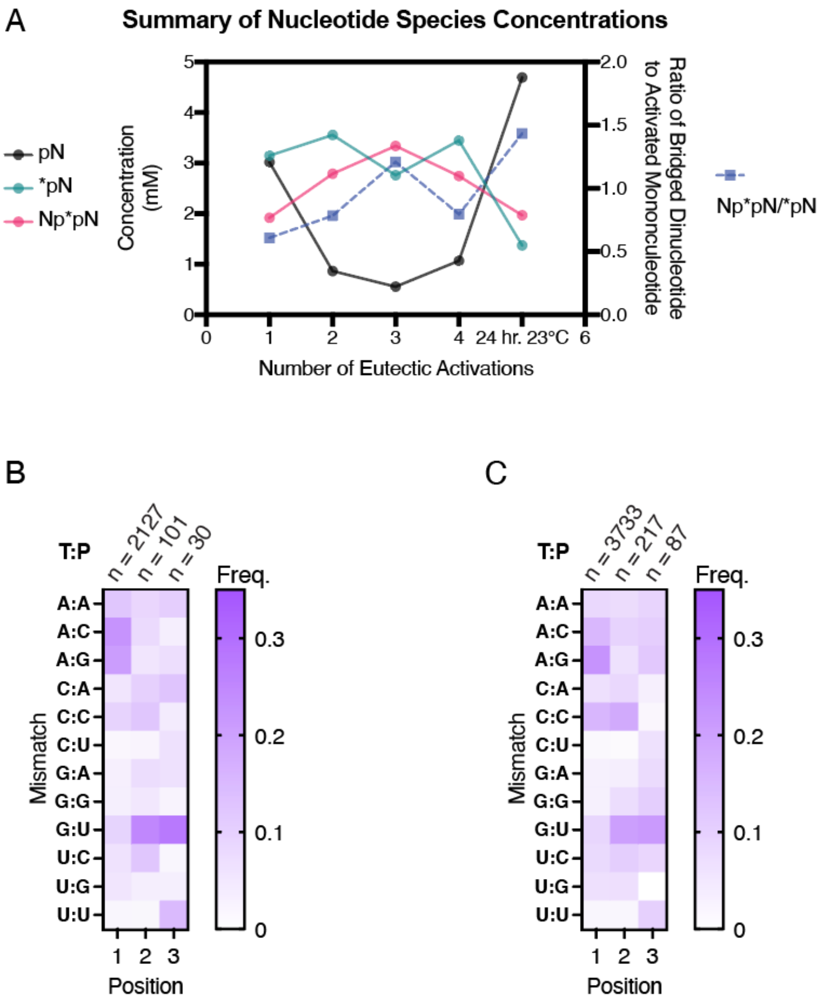


**Figure S8.** Eutectic phase activation maintains a high ratio of bridged dinucleotides to activated mononucleotides. (A) The concentrations of nucleotide species during eutectic phase activation and a subsequent 24-hour incubation with bridge-forming chemistry of all four nucleotides based on NMR data.

*Reaction conditions:* 2.5 mM each rNMP (10 mM total), 30 mM $MgCl_2$, 50 mM $Na^+$-HEPES, pH 8, 100 mM 2MBA, and 50 mM MeNC were subjected to four 24 hour rounds of eutectic phase activation with the addition of 50 mM MeNC at each thaw. The subsequent 24 hour incubation was at 23°C. Bridge-forming chemistry was the addition of 100 mM 2MBA and 200 mM MeNC. (B) The position-dependent frequency of each mismatch, relative to all mismatches (n) at the indicated position from the same experiment used to generate Figure S6C. (Activated nucleotides generated by four rounds of eutectic activation were added to the sequencing hairpin and incubated for 24 hours with bridge-forming chemistry.) T:P = Template:Product. (C) The position-dependent frequency of each mismatch, relative to all mismatches (n) at the indicated position from the same experiment used to generate Figure 5B and S6D. (The sequencing hairpin was included during the four rounds of eutectic activation, and then incubated for 24 hours with bridge forming chemistry.) T:P = Template:Product.

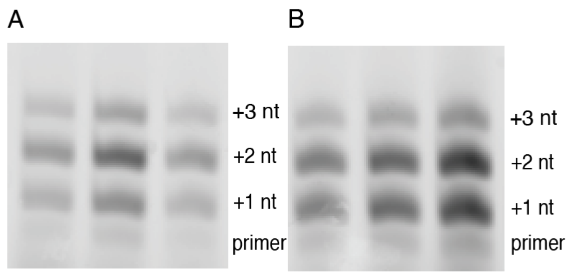


**Figure S9.** Eutectic ice phase completes a pathway from unactivated mononucleotides through activation to primer extension. Replicates of primer extension reactions using (A) pure activated mononucleotides (5 mM *pG and 5 mM *pC) under room temperature and (B) pNs (pN = pC and pG) generated with isocyanide activation chemistry under ice eutectic phase. Positions of primer and +1 to +3-nucleotide (nt) extension products are indicated.
*Reaction conditions:* (A) 5 mM *pG and 5 mM *pC were added to 1 μM primer, 1.5 μM template, 200 mM $Na^+$-HEPES pH 8.0, and 30 mM $MgCl_2$. (B) 1 μM primer and 1.5 μM template were added to a mixture of 10 mM pN, 10 mM 2AI, 100 mM 2MBA, 30 mM $MgCl_2$, 50 mM $Na^+$-HEPES pH 8, after four cycles of eutectic phase activation (50 mM MeNC each addition).

**Table S1. Numerical Data Used to Generate Heat-map in Figure 5C**

Frequency

| Product Base | | | |
|---|---|---|---|
| **A** | **0.141** | **0.052** | **0.036** |
| **C** | **0.610** | **0.537** | **0.720** |
| **G** | **0.226** | **0.405** | **0.210** |
| **U** | **0.023** | **0.006** | **0.033** |
| | **1** | **2** | **3** |

Position

**Table S2. Numerical Data Used to Generate Heat-map in Figure 5D**

Frequency

| Bridged Dinucleotide | | | |
|---|---|---|---|
| U*U | 0.001643 | 0.000294 | 0.004324 |
| U*G | 0.009573 | 0.001057 | 0.010687 |
| U*C | 0.008852 | 0.003107 | 0.018134 |
| U*A | 0.004070 | 0.002055 | 0.004206 |
| G*U | 0.030122 | 0.080182 | 0.064413 |
| G*G | 0.074462 | 0.105564 | 0.039132 |
| G*C | 0.078178 | 0.165004 | 0.080551 |
| G*A | 0.030034 | 0.034081 | 0.013471 |
| C*U | 0.088771 | 0.093670 | 0.095016 |
| C*G | 0.158480 | 0.088228 | 0.136500 |
| C*C | 0.192083 | 0.192342 | 0.389887 |
| C*A | 0.171804 | 0.178669 | 0.106839 |
| A*U | 0.005688 | 0.002214 | 0.005282 |
| A*G | 0.076777 | 0.027086 | 0.010852 |
| A*C | 0.051003 | 0.020643 | 0.020706 |
| A*A | 0.018460 | 0.005803 | 0.000000 |
| | 1 | 2 | 3 |

Position

**Table S3. Numerical Data Used to Generate Heat-map in Figure S8B**

Frequency

| Mismatch (Template:Product) | | | |
|---|---|---|---|
| A:A | 0.1210 | 0.0854 | 0.1027 |
| A:C | 0.2225 | 0.0776 | 0.0342 |
| A:G | 0.2051 | 0.0543 | 0.0685 |
| C:A | 0.0537 | 0.0985 | 0.1257 |
| C:C | 0.0925 | 0.1204 | 0.0419 |
| C:U | 0.0154 | 0.0219 | 0.0629 |
| G:A | 0.0315 | 0.0718 | 0.0632 |
| G:G | 0.0339 | 0.0497 | 0.0253 |
| G:U | 0.0913 | 0.2487 | 0.2782 |
| U:C | 0.0613 | 0.1216 | 0.0165 |
| U:G | 0.0538 | 0.0358 | 0.0329 |
| U:U | 0.0181 | 0.0143 | 0.1481 |
| | 1 | 2 | 3 |

Position

**Table S4. Numerical Data Used to Generate Heat-map in Figure S8C**

Frequency

Mismatch (Template:Product)

| | | | |
|---|---|---|---|
| **A:A** | **0.0820** | **0.0735** | **0.0906** |
| **A:C** | **0.1520** | **0.0931** | **0.1035** |
| **A:G** | **0.2244** | **0.0637** | **0.1165** |
| **C:A** | **0.0656** | **0.0829** | **0.0317** |
| **C:C** | **0.1540** | **0.1796** | **0.0158** |
| **C:U** | **0.0123** | **0.0069** | **0.0634** |
| **G:A** | **0.0302** | **0.0349** | **0.0765** |
| **G:G** | **0.0302** | **0.0733** | **0.1052** |
| **G:U** | **0.0880** | **0.2024** | **0.2103** |
| **U:C** | **0.0789** | **0.1039** | **0.0871** |
| **U:G** | **0.0660** | **0.0677** | **0.0000** |
| **U:U** | **0.0164** | **0.0181** | **0.0995** |
| | **1** | **2** | **3** |

Position
